# Finding functional disease-associated non-coding variation using next-generation sequencing

**DOI:** 10.1101/060285

**Authors:** Paolo Devanna, Xiaowei Sylvia Chen, Joses Ho, Dario Gajewski, Alessandro Gialluisi, Clyde Francks, Simon E. Fisher, Dianne Newbury, Sonja C. Vernes

## Abstract

Next generation sequencing has opened the way for the large scale interrogation of cohorts at the whole exome, or whole genome level. Currently, the field largely focuses on potential disease causing variants that fall within coding sequences and that are predicted to cause protein sequence changes, generally discarding non-coding variants. However non-coding DNA makes up ^~^98% of the genome and contains a range of sequences essential for controlling the expression of protein coding genes. Thus, potentially causative non-coding variation is currently being overlooked. To address this, we have designed an approach to assess variation in one class of non-coding regulatory DNA; the 3′UTRome. Variants in the 3'UTR region of genes are of particular interest because 3'UTRs are responsible for modulating protein expression levels via their interactions with microRNAs. Furthermore they are amenable to large scale analysis as 3′UTR-microRNA interactions are based on complementary base pairing and as such can be predicted *in silico* at the genome-wide level. We report a strategy for identifying and functionally testing variants in microRNA binding sites within the 3'UTRome and demonstrate the efficacy of this pipeline in a cohort of language impaired children. Using whole exome sequence data from 43 probands, we extracted variants that lay within 3'UTR microRNA binding sites. We identified a common variant (SNP) in a microRNA binding site and found this SNP to be associated with an endophenotype of language impairment (non-word repetition). We showed that this variant disrupted microRNA regulation in cells and was linked to altered gene expression in the brain, suggesting it may represent a risk factor contributing to SLI. This work demonstrates that biologically relevant variants are currently being under-investigated despite the wealth of next-generation sequencing data available and presents a simple strategy for interrogating non-coding regions of the genome. We propose that this strategy should be routinely applied to whole exome and whole genome sequence data in order to broaden our understanding of how non-coding genetic variation underlies complex phenotypes such as neurodevelopmental disorders.

## INTRODUCTION

Next generation sequencing (NGS) is a powerful approach for identifying genetic variation contributing to human traits and has proven to be particularly valuable for uncovering rare variation underlying complex disorders. Whole exome sequencing (WES) has been widely applied to this problem as it represents an affordable survey of the gene-coding portion of the human genome, which accounts for about 1% of the total sequence. WES has recently been used to reveal new candidate genes and pathways for neurodevelopmental disorders such as language impairment, intellectual disability and autism (1–4), which have previously been largely intractable.

The greatest strength of NGS, the ability to rapidly obtain genome-wide sequence information, also represents one of its greatest challenges. Inter-individual differences at the sequence level mean that WES identifies >20,000 variants per person (5), and this number jumps to a staggering >3 million variants when whole genome sequencing (WGS) is performed (6). Understanding which variants are related to the phenotypes under study out of the large number of variants identified requires accurate analysis and robust biological interpretation of the data. In order to make sense of these data it is necessary to prioritise some variants and filter out those changes that are considered less likely to contribute to the phenotype or disorder being studied. In practice, this routinely involves the removal of variants that are common in the general population, that are found outside protein coding regions, or that do not directly affect protein sequence. Thus, non-coding variants that may be located in intergenic regulatory elements are often being discarded, despite their potential functionality and relevance to phenotypes. Regulatory changes are of particular interest given that precise control of gene expression is essential to normal development and function and underlies normal brain circuit formation and ongoing neuronal activity. We posit that it is possible to understand the contribution of both rare and common intergenic variants if existing data characterising non-coding DNA (e.g. coordinates for regulatory regions, histone marks etc.) are routinely combined with NGS data during analysis pipelines. Only in this way will a truly genome-wide picture of the genetic factors (both coding AND non-coding) that underlie complex neurodevelopmental disorders be built. To this end, we designed an approach for using NGS data to identify and importantly to also functionally test the effects of variation in one class of non-coding DNA; the 3′UTRome. Herein we show the efficacy of this method using WES data from a cohort of children classified as having a Specific Language Impairment (SLI).

The 3 prime untranslated regions (3′UTR) of genes have long been considered likely candidates for pathogenic mutations (7–10) and have been implicated in a range of neurological disorders. Sequencing of the 3′UTRs of genes involved in tinnitus, Parkinson′s disease, Tourette′s syndrome and autism have linked common SNPs and rare non-coding variants to these disorders (9–11). *SLITRK1* has been linked to Tourette′s syndrome and in a screen searching for further variants in this gene, a 3′UTR variant was identified in two unrelated Tourette′s patients (this variant was absent from >4000 control samples) (12). The variant fell within a microRNA (miR) binding site (miR-189) of the *SLITRK1* 3′UTR. Functional studies showed that miR-189 represses *SLITRK1* expression by interacting with the control allele in the 3'UTR binding site. Strikingly, the patient-identified variant significantly altered this regulation, suggesting a functional link between this non-coding variant and Tourette′s syndrome (12). In addition to evidence linking 3'UTRs to disorders, these regions are strong candidates for study for five key reasons; (i) 3′UTR regions are found at the end of all protein coding genes in the genome and are comprehensively annotated (13); (ii) 3′UTR regions directly affect the amount of protein produced from a gene via interactions with small regulatory RNA molecules known as microRNAs; (iii) microRNA interactions are based on complementary nucleotide pairing with short sequences (7-20 bp) in the 3′UTR regions and thus can be accurately predicted from sequence data; (iv) WGS captures all variation in the genome, including the 3'UTRome, and thus all functional variation in this region can be identified. WES platforms are designed to capture only the gene-coding exonic sequences, however they also capture some flanking DNA regions, meaning that a reasonably large amount of 3′UTR sequence data can also be extracted from the wealth of existing and ongoing WES data available and; (v) functional effects of observed variants can be tested in the lab using simple, scalable assays.

Specific language impairment (SLI), the failure to acquire age appropriate language skills in the absence of any explanatory factors (e.g. learning difficulties), affects up to 8% of school age children (14, 15). Despite strong evidence for genetic underpinnings of language impairment, efforts to identify causative factors via linkage and association studies have found multiple different genomic regions, all predicted to have small effects (16–18). Thus, like many other neurodevelopmental disorders, SLI seems to have a heterogeneous set of genetic risk factors contributing to the phenotype. As such, an effort was made to identify causative genetic factors via exome sequencing of a cohort of children with severe SLI (Chen et al, this issue). This study interrogated the protein coding regions of the genome and, like the overwhelming majority of exome sequencing studies, focused largely on variation that was predicted to alter protein sequence. This is a widespread strategy when attempting to identify likely pathogenic variants as *in silico* prediction programs principally focus on protein coding changes. Non-coding changes are less well studied and thus often classified by prediction programs as having uncertain/unknown function. Using WES data, we demonstrate herein that it is possible to identify putative causative variants in non-coding regions and importantly that these variants can have direct functional consequences, making them strong candidates for pathogenicity.

## RESULTS

### 3'UTR variants within microRNA binding sites can be identified from WES data

We designed a pipeline to identify and assess the functionality of both common and rare single nucleotide variants (SNVs) identified in non-coding 3′UTR regions of the genome and applied this to a WES dataset from a cohort of 43 children with severe SLI (Chen et al, this issue) (Figure 1).

Chen et al (this issue) identified all SNVs present in the WES data throughout the genome. To rule out likely false positives, only SNVs with >10× sequence coverage were retained (see Supplementary Table S1 for SNV numbers identified in different regions of the genome in this dataset). From all filtered variants we extracted only those SNVs that were within the 3′UTR region of a gene (N = 6606, over 4651 3′UTR regions). 3′UTRs can be as short as 60 bp and in some extreme cases as long as 20 kilobase pairs (kb) (19), but in the human genome they are composed of approximately 800 bp on average (20). In the SLI WES dataset, for the majority of 3'UTRs in the genome (62.7% of 3′UTRs annotated by Ref-Seq), the first ^~^200 bp had a read depth ≥10 allowing the reliable identification of variants in this region (Figure 2A). The highest density of microRNA binding sites is also found in this initial region of the 3′UTR (Figure 2B), suggesting that a large number of binding sites can be assessed per gene, even using the limited region of the 3'UTR covered by standard exome sequencing.

**Figure 1.**
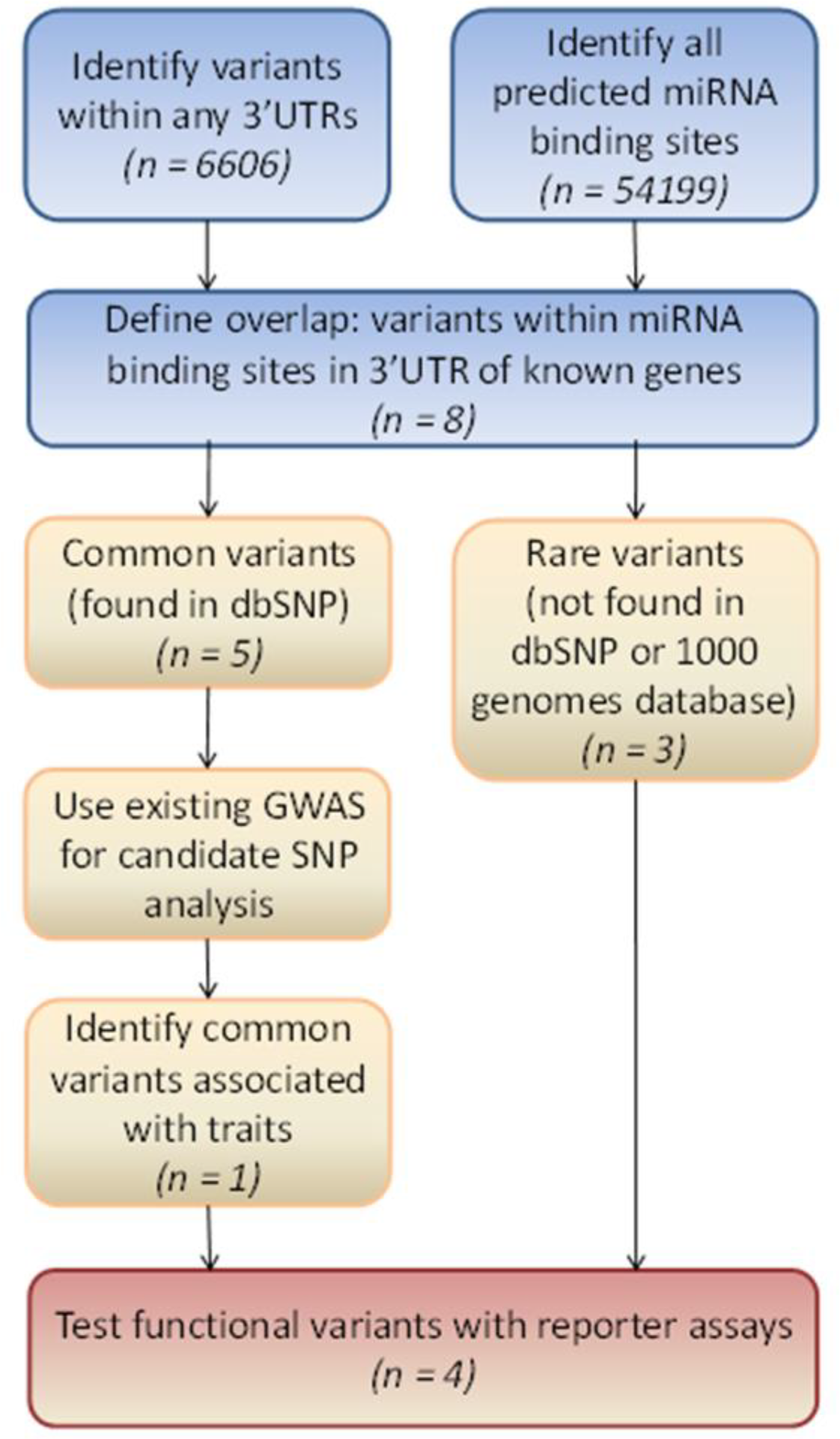
Workflow for identification of non-coding 3'UTRome variants in next generation sequencing (NGS) data. This flowchart demonstrates how variants can be identified in non-coding regulatory regions of the genome using NGS data. First, a list of all variants found within the 3'UTR regions of any genes was generated from WES data (N = 6606). This was overlapped with all predicted microRNA binding sites in the genome (via targetscan 6.2) (N = 54199) to create a list of variants that lie within miRNA binding sites in the 3'UTR of known genes (N = 8). The identified variants fell into two categories; 3 rare variants found in a single proband and 5 common variants that were annotated in dbSNP. Common variants were further assessed using candidate SNP analysis in a GWAS study of a large SLI cohort. One of the common variants showed association to a quantitative measure of language impairment. SNVs that pass the bioinformatic screening (N = 4) were characterized using reporter assays to demonstrate functionality of the wild type site and consequences of patient identified variants. For each stage the number of variants/sites identified in each category in our study is shown in brackets.

**Figure 2.**
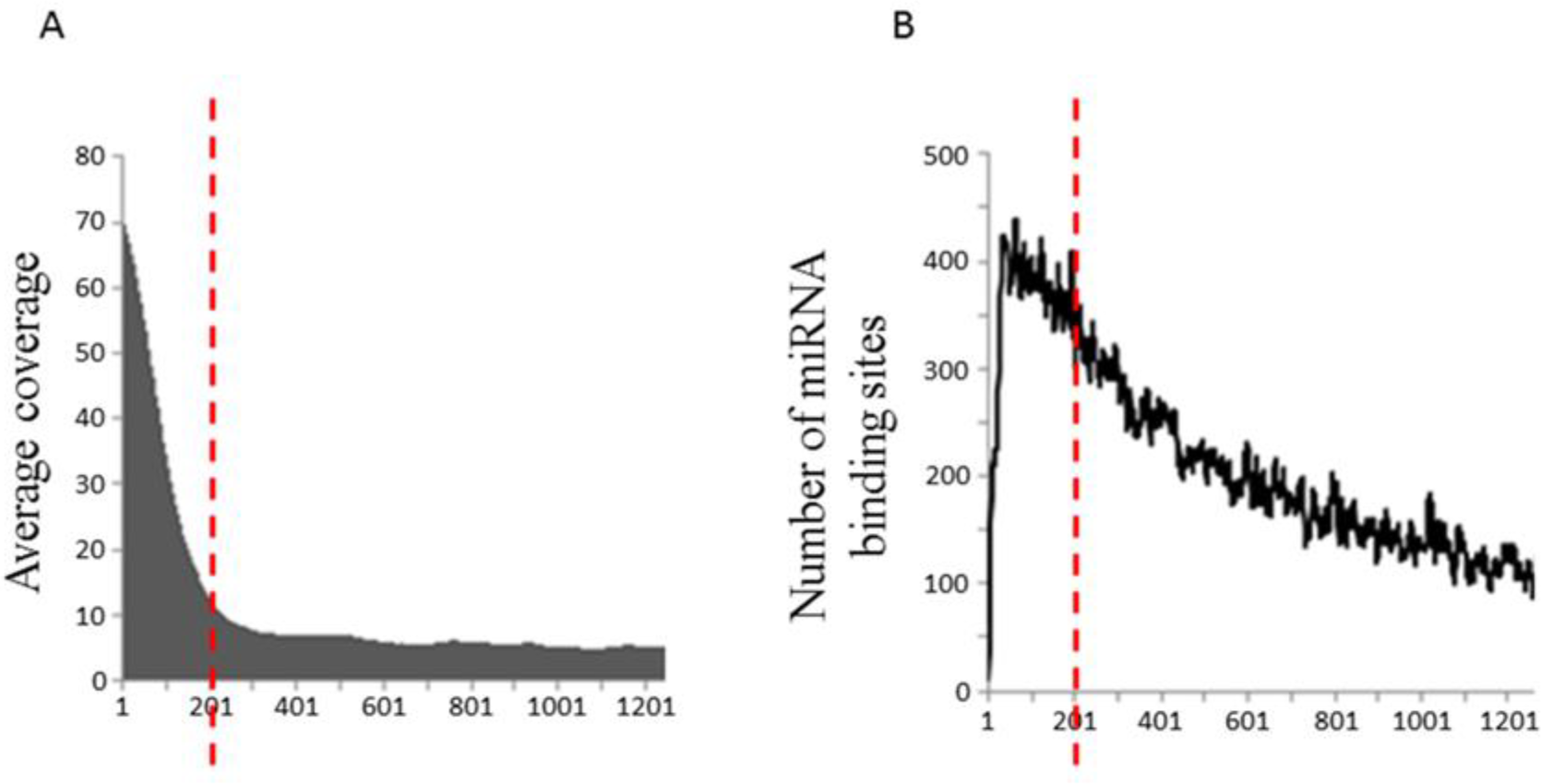
Identification of non-coding variants in exome sequencing data. **(A)** Average read depth profile across the 3′UTRome in the SLI WES dataset **(B)** Distribution of predicted microRNA binding sites across the human 3′UTRome (predicted by Targetscan). The red dotted lines indicate the 200bp boundary.

To determine if any of the 3'UTR variants fell within microRNA binding sites, we separately searched for all predicted sites using the TargetScan algorithm (21) and overlaid these with the WES identified SNVs. Eight 3′UTR SNVs were thus identified within a sequence predicted to be bound by a microRNA. Three of these were rare variants, only found in a single individual and each in the 3′UTR of different genes; *BTN2A1*, *CENPJ* and *MTMR3* (Table 1). These were considered to be private mutations as they were not present in other datasets (dbSNP (22), 1000 genomes (23), ExAc browser (24)), indicating that they have a population frequency of less than 0.00082%. The presence of these rare variants in each of the relevant probands was confirmed with bi-directional Sanger sequencing (Figure S1). Five of these SNVs were identified from dbSNP as single nucleotide polymorphisms (SNPs) found in the general population (frequency >0.1%); four are common (population frequency > 1%), and one is rare (0.3%) (Table 1).

**Table 1.** SNVs located within predicted microRNA binding sites in SLI probands.

| Chr | Position | dbSNP | dbSNP<br>Global<br>MAF<br>freq | Ref | Alt | Proba<br>nds | Gene | microRNA |
| --- | --- | --- | --- | --- | --- | --- | --- | --- |
| 6 | 26469054 | - | - | G | C* | 1 | BTN2A1 | miR-342 |
| 13 | 25457299 | - | - | G | T* | 1 | CENPJ | miR-433 |
| 22 | 30421860 | - | - | C | T* | 1 | MTMR3 | miR-346 |
| 9 | 35661943 | rs72727021 | 0.0469 | A | C* | 9 | ARHGEF39 | miR-192/215 |
| 16 | 87436764 | rs1054528 | NA | A* | G | 42 | MAP1LC3B | miR-204/211 |
| 16 | 79245820 | rs383362 | 0.3223 | G | T* | 34 | WWOX | miR-134/758 |
| 19 | 6751293 | rs1049232 | 0.2404 | T | G* | 13 | TRIP10 | miR-17-5p |
| 19 | 53086946 | rs190191374 | 0.0030 | G | T* | 1 | ZNF701 | miR-199/199-5p |

\*minor allele

### A common variant within a microRNA binding site is associated with SLI

To determine the relevance of the common (SNP) variants to SLI we took advantage of genome-wide SNP data from the SLI Consortium (SLIC) sample. This dataset comprised 285 SLI families recruited from five centres across the UK; The Newcomen Centre at Guy′s Hospital, London (now called Evelina Children′s Hospital); the Cambridge Language and Speech Project (CLASP); the Child Life and Health Department at the University of Edinburgh; the Department of Child Health at the University of Aberdeen; and the Manchester Language Study (16, 18). The 43 probands sequenced herein are a subset of this cohort. Two common SNPs (rs1054528 and rs190191374) were not genotyped in the SLIC (SLI Consortium) dataset and thus could not be assessed. However, three of the five common SNPs were directly genotyped or imputed from the SLIC dataset (rs72727021 (imputed), rs383362 (genotyped) and rs1049232 (genotyped)) and thus we could use these data for a candidate association analysis. Genotype data were available for 983 parents and children from these 285 SLI families. As previously described, various quantitative measures of language-related abilities were available for children in this cohort (16, 25, 26). Measures of expressive (ELS) and receptive (RLS) language were obtained using the Clinical Evaluations of Language Fundamentals (CELF-R) (27). Verbal IQ (VIQ) and performance IQ (PIQ) were assessed using the Wechsler Intelligence Scale for Children (28) and reading and spelling ability were measured with the Wechsler Objective Reading Dimensions (WORD)(29). In addition, the phonological short-term memory of adults and children was assessed with a 28-item test of non-word repetition (NWR) (30). Association was assessed within and between family units using the QFAM test in PLINK. This quantitative test of association employs an adaptive permutation procedure to account for the dependence between related individuals. One SNP (rs72727021), located in the 3'UTR of the *ARHGEF39* gene (also known as *C9orf100*), was marginally associated with the non-word repetition measure in the SLI cohort (empirical p= 7.7 × 10^-3^) (Supplementary Table S2). The alternative allele (C) of rs72727021, which has a population frequency of 4.7% (10.8% in 120 CEPH individuals in 1000 genomes pilot phase I) is carried by 23% of the SLIC probands (MAF=12.3%) and is associated with a reduction of 10 points (0.66SD) on the NWR test in these individuals.

### Functional validation of a WES 3'UTR variant located within a microRNA binding site

From our exome data we have identified four candidate variants located in microRNA binding sites that may be related to the SLI phenotype; one common variant (rs72727021) that is associated with NWR in the SLI cohort, and three rare (private) variants. Because these microRNA binding sites were identified via *in silico* predictions, we first set out to functionally validate the predicted interaction between these 3'UTR binding sites and their cognate microRNAs using a reporter assay. We cloned an approx. 300-400 bp region from each genes 3'UTR, spanning the predicted miRNA binding site (Figure 3A). This was inserted into an expression vector downstream of the luciferase reporter gene as part of its 3'UTR. We then determined the ability of the relevant microRNAs to regulate the predicted binding site within the 3'UTR of each gene. The binding sites identified within the 3'UTRs of *BTN2A1, CENPJ* and *MTMR3* were not regulated by their predicted microRNAs (miR-342, miR-433 and miR-346 respectively) in these reporter assays, suggesting they are not functional binding sites (Figure 3B). As such we conclude that the rare variants found within these binding sites are unlikely to disrupt microRNA-3′UTR interactions, and thus unlikely to directly contribute to SLI via this regulatory mechanism. However, 3'UTRs can have other functions, and as such we cannot rule out that the presence of these variants might affect other 3'UTR dependent post-transcriptional processes and lead to SLI related phenotypic outcomes.

The binding site within the 3'UTR of *ARHGEF39* is predicted to be recognised and bound by both miR-215 and miR-192. The seed sequence for both these microRNAs is identical and both microRNAs have the same confidence score (as predicted by Targetscan), suggesting that they are equally able to bind this site. Given this equivalence we used miR-215 to demonstrate the functionality of this site. The *ARHGEF39* 3'UTR reporter containing the reference allele of the rs72727021 SNP was significantly downregulated by miR-215 (Figure 3B). This demonstrates that the microRNA binding site found in the 3'UTR of *ARHGEF39* is functional and that miR-215 downregulates expression by interacting with this site when the reference allele ('A') is present.

We then went on to determine how the presence of the SLI-associated alternative allele ('C') affected this regulatory relationship. Introducing the alternative 'C' allele into the 3'UTR of the reporter construct abolished regulation by miR-215 (Figure 3C). To confirm the specificity of this interaction we also made a variant where the miR-215 binding site in the 3′UTR was completely deleted (DEL). As expected miR-215 could no longer regulate the reporter construct when the deletion was introduced (Figure 3C). Interestingly this loss of regulation was not significantly different to the control vector (lacking a 3'UTR) or the vector carrying the alternative 'C' allele of rs72727021 in its 3'UTR.

Together this demonstrates that the *ARHGEF39* 3'UTR can be down-regulated by miR-215 when the rs72727021 reference allele ‘A′ is present. If the alternative 'C' allele is present, miR-215 regulation is abolished leading to higher gene expression, and this effect is as severe as a complete deletion of the miR-215 binding site.

**Figure 3.**
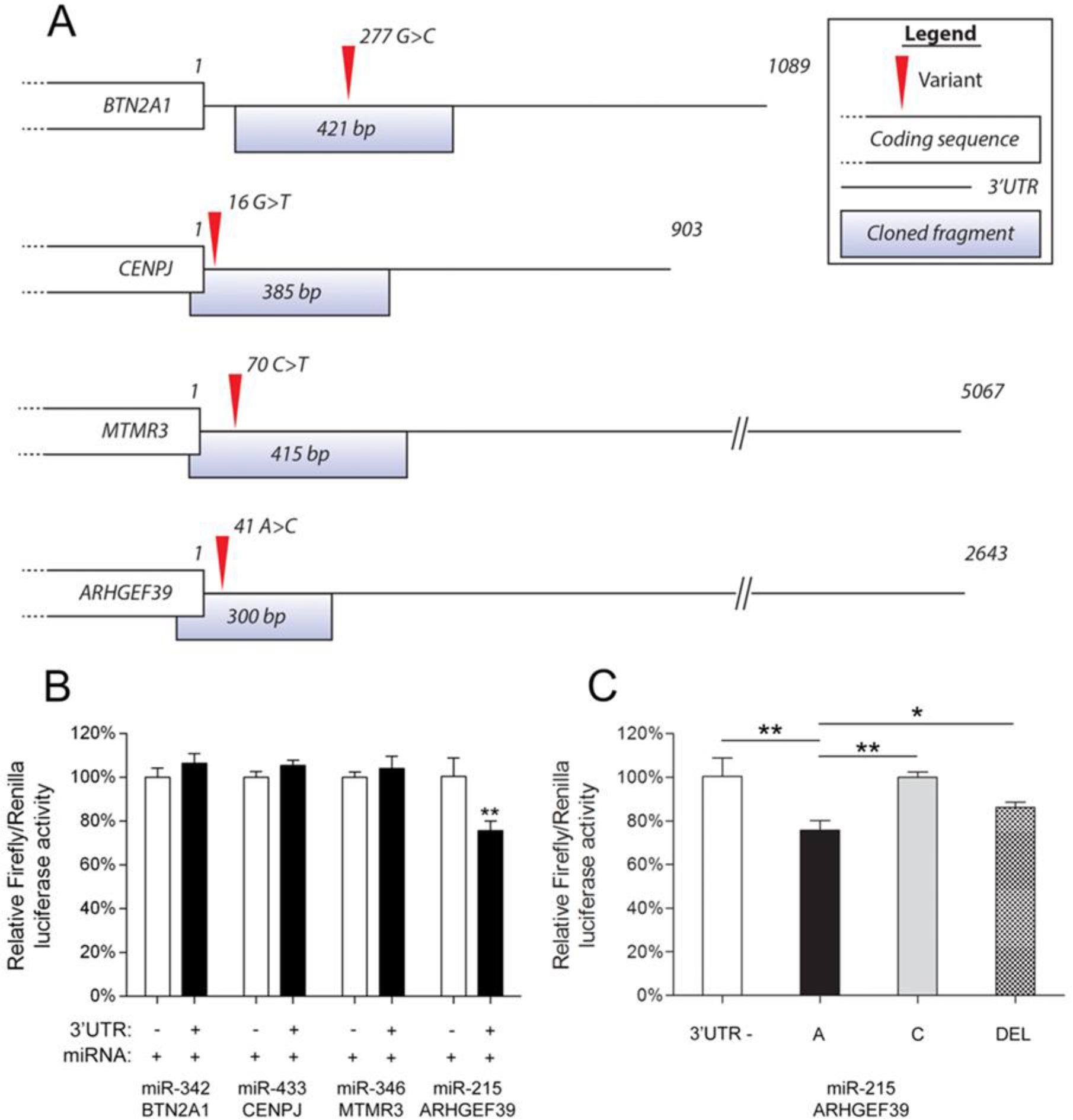
Functional consequences of 3′UTR variants identified in SLI exomes. (A) 3′UTR fragments of *BTN2A1*, *CENPJ*, *MTMR3* and *ARHGEF39* that spanned the predicted microRNA binding sites were cloned downstream of the bioluminescent luciferase reporter gene. This schematic illustrates the length of the 3'UTR (depicted by a straight line) relative to the end of the protein coding sequence of each gene (depicted by a white box), the cloned fragments that was used in reporter assays, and the identity and position of the 3'UTR variants identified in the SLI cohort. (B) Luciferase reporter assays were used to demonstrate if predicted binding sites were functionally regulated by the predicted microRNAs. Candidate 3'UTR regions (shown in part A) were cloned downstream of the luciferase reporter gene. These UTR reporter constructs (3'UTR: +) were co-expressed with the microRNA that was predicted to regulate each UTR. To determine specificity of this regulation, control reporters were also tested that lacked any cloned fragment (3'UTR: -). Expression of the reporter gene was the same with (+) or without (-) the 3'UTR fragment for *BTN2A1, CENPJ* and *MTMR3*, showing that these sites were not regulated by these microRNAs (miR-342, miR-433 and miR-346). In contrast, reporter gene expression was significantly lower when the *ARHGEF39* 3'UTR fragment was present (+) compared to without any 3'UTR (-), showing that miR-215 represses gene expression by interacting with the 3'UTR when the reference allele 'A' is present. No differences in reporter activity were observed in absence of miR-215 co-expression (data not shown) (C) To determine if the presence of the alternative 'C' allele disrupts this regulatory relationship, we introduced the 'C' allele into the reporter gene UTR. The presence of the SLI-associated 'C' allele abolished repression of the reporter gene by miR-215, showing its biological relevance. To show specificity of this effect the entire miR-215 binding site was deleted from the *ARHGEF39* 3'UTR reporter ('DEL') and this construct was also not regulated by miR-215. Deleting the entire miR-215 binding site ('DEL') was not significantly different to the effect of introducing the 'C' allele and neither of these were significantly different from the construct that had no 3'UTR cloned fragment present ('3'UTR -'). Significant differences between groups were calculated using an ANOVA test followed by post-hoc Tukey calculation. Only statistically significant differences are noted in the figure. Significance is indicated by *p < 0.05 and ** p < 0.01. All results are reported as the average +/− standard deviation of 3 biological replicates.

### *In vivo* brain expression differences are associated with rs72727021

In addition to these functional assays, we also identified strong support for the relevance of this variant in controlling *ARHGEF39* expression levels *in vivo*. Using eQTL data from the GTEx portal (31) we found that that the rs72727021 SNP was significantly associated with *ARHGEF39* expression in the human brain. Cortical samples from individuals heterozygous or homozygous for the alternative ‘C′ allele had higher *ARHGEF39* expression (cortex p=7.5e-8; frontal cortex p=5.1e-9) (Figure 4). This data mirrors the loss of repression (higher reporter gene expression) we observed when the alternative 'C' allele was present (compared to the reference 'A' allele) during *in vitro* functional assays (Figure 3C). To support the specificity of this *in vivo* data we checked the GTEx portal for eQTL effects related to the other 4 common variants identified in this cohort (Table 1). No significant differences in expression were observed for these variants in the human brain, further supporting the significance of our findings.

**Figure 4.**
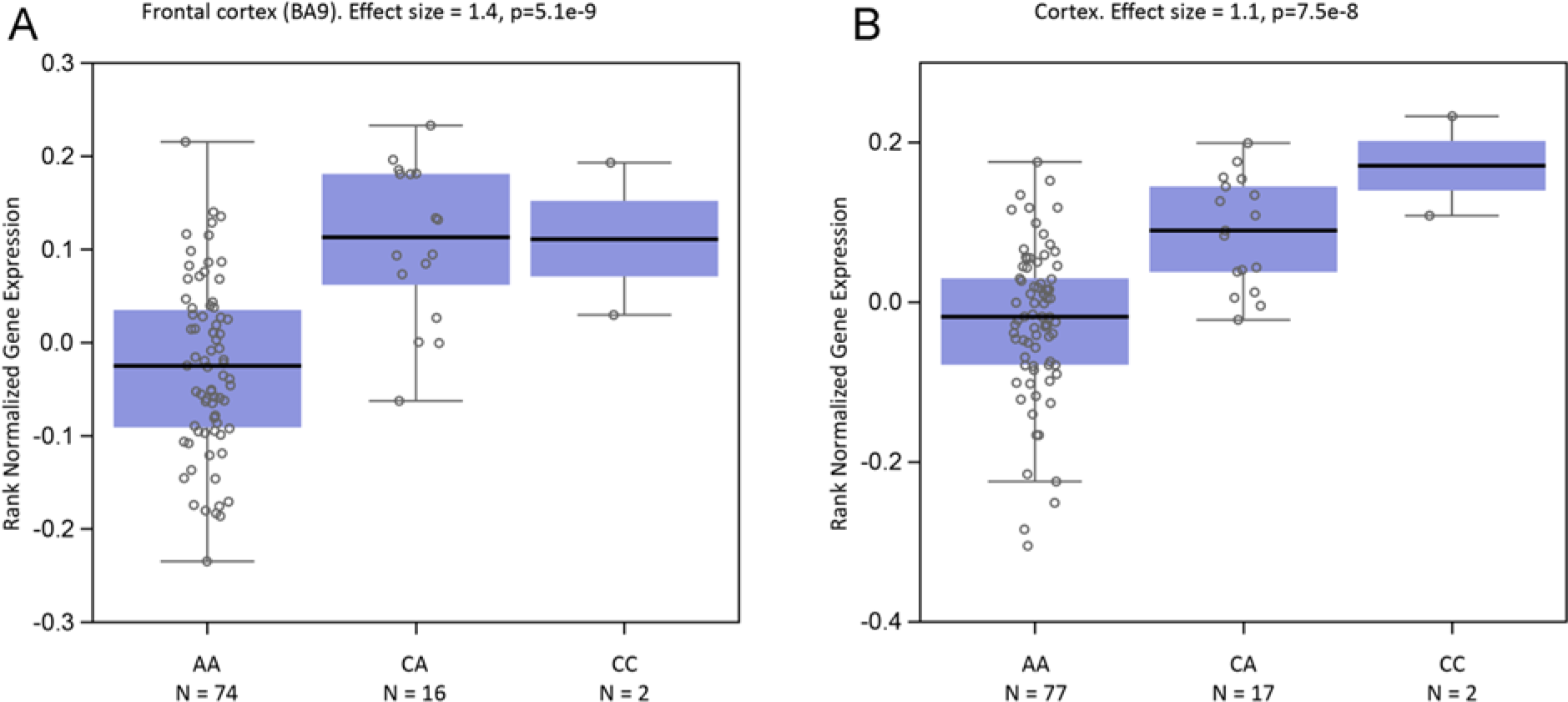
rs72727021 is associated with ARHGEF39 expression in the human brain. eQTL data was obtained from the GTEx database. Significant associations were observed between rs72727021 and *ARHGEF39* expression in samples from (A) the frontal cortex (BA9) (effect size = 1.4, p=5.1e-9) and (B) the cortex (effect size = 1.1, p=7.5e-8). In both sample sets ARHGEF39 expression is higher when the rs72727021 alternative allele ('C') is present in the heterozygous or homozygous state.

Taken together, these findings show that we have identified a common non-coding SNP with a significant association to SLI and a functional effect on gene expression in both cell models and the human brain. These data represent the first time an SLI-associated SNP has been reported to have direct functional consequences.

### rs72727021 in other datasets with language related metrics

Given the novelty of this finding and the functional effect of this SNP *in vitro* and *in vivo*, we sought to find supportive evidence for the association between rs72727021 and NWR in other datasets. To our knowledge a true replication dataset does not exist, in the sense of having both a similar recruitment scheme and the same measurement of NWR as the SLIC dataset. We nonetheless queried the association in two independent datasets; the Avon Longitudinal Study of Parents and Children (ALSPAC) cohort (32) and the Colorado Learning Disabilities Research Centre (CLDRC) dataset (33). The ALSPAC cohort included 1681 individuals that were recently sequenced at low coverage as part of the UK10K project (34) and underwent a short (12-item) form of the non-word repetition test. Ethical approval for the study was obtained from the ALSPAC Ethics and Law Committee and the Local Research Ethics Committees. All data relating to the ALSPAC study was collected by the ALSPAC team. The ALSPAC study website contains details of all the data that is available through a fully searchable data dictionary and reference the following webpage http://www.bris.ac.uk/alspac/researchers/data-access/data-dictionary/. The CLDRC dataset included 727 children of which 701 subjects were tested for NWR (mean age 11.7 years, age range 8-19), composed by twins recruited in Colorado (US) for a school history of RD or ADHD, along with their co-siblings (343 unrelated twinships/sibships) (33). Quantitative association analyses were performed within PLINK, but no significant association was found in these datasets. It is important to note that the lack of support from these cohorts may reflect variations in sample ascertainment since the SLIC population was specifically selected for severe language phenotypes, or subtle differences in the tests used to generate the metrics related to language ability. Further investigations in specifically selected samples will be needed to clarify the genetic association between rs72727021 and language phenotypes.

### DISCUSSION

Herein we demonstrate an approach for identifying and validating the functionality of non-coding variants from NGS data. Given the importance of non-coding DNA for regulating gene expression and the simple approach we outline for assessing these variants, we strongly recommend that such 3'UTRome variation be considered as standard in all NGS pipelines.

Although current WES platforms do not sequence the entire 3'UTR (by design), we have shown that we can extract functional variants from exome sequencing. The widespread availability of WES data makes it a cost-effective and practical way to discover functionally relevant SNVs related to complex phenotypes. To date, more than 1 million exomes have been sequenced using WES and the methodology we propose here is immediately applicable to any existing WES datasets for which raw sequence reads or called variants are available. Methods that interrogate greater portions of the genome, such as whole genome sequencing will give a wealth of information about the complete 3'UTRome and make our pipeline even more valuable for its ability to discover functional non-coding variation. Studies of complex disorders are increasingly making use of WGS, and including our pipeline as standard when assessing WGS data from large cohorts will allow unprecedented identification of functional, non-coding risk factors. A similar approach could also be used to identify potentially functional common variants from GWAS data, although the coverage of microRNA binding sites will be substantially lower in such datasets.

A major strength of this approach is that it provides a simple, high throughput method for validating the functional consequences of non-coding 3'UTRome variants. Although pathogenicity prediction algorithms are widely used, the high false positive/negative rates (35, 36) for both coding and noncoding variation reinforces the importance for functional testing in pipelines designed to link genetic variation to phenotypes. Functional validation is possible for protein coding variants, however it may not always be feasible given the wide range of tests required to explore the diverse functions of different protein classes. Only a small proportion of studies have shown protein-level effects of WES/WGS identified variants. Those that have often focus on basic protein features such as protein stability and subcellular localization due to practical limitations, (37-39) potentially resulting in a high false negative rate. Because the functionality of a microRNA-3′UTR interaction can be determined via a simple set of reporter gene assays, this approach facilitates high throughput and reliable assessment of the identified variants. Furthermore, our method has the advantage of requiring minimal bioinformatic analysis (see materials and methods) if called variants are available, making it an attractive approach to bench scientists.

This work also represents the first report of functional consequences for a non-coding SNP associated with SLI and highlights a possible role for the rs72727021 variant and/or the *ARHGEF39* gene (in which the variant is found) in language disorder. Little is known regarding the *ARHGEF39* gene, but it is a member of the ARHGEF family of Rho guanine nucleotide exchange factors. These proteins act as molecular switches to regulate diverse processes including transcriptional regulation, cell migration, cell growth and dendritic development (40-42). Other members of this family have previously been implicated in language impairment (*ARHGEF19*) (16) and intellectual disability (*ARHGEF6*) (43). Future studies into the expression pattern and function of this gene may reveal a potential role for *ARHGEF39* in the development of language relevant circuitry in the brain.

In summary, we report an accessible, rapid and biologically relevant method for assessing noncoding variation that adds significantly to our ability to identify causative variants from WES and WGS data – a major challenge facing the future of genomics. Adding this simple pipeline to the standard WES/WGS toolkit is expected to reveal a wealth of functional variation in previously overlooked non-coding regions of the genome and will help to identify new links between genes and complex disorders.

## MATERIALS AND METHODS

### Variant identification

Full methods on WES sequencing and variant calling are described by Chen et al (this issue). Briefly, for sequencing, the exome was captured by SureSelect Human All Exon version-2 50 Mb kit (Agilent, Santa Clara, CA, USA), sequenced using the SOLiD series 5500xl DNA sequencing platform (Life Technologies, Carlsbad, CA, USA) and called via the standard BWA-GATK pipeline, followed by quality filtering as described by Chen *et al*.

Genome wide coordinates (hg19) for microRNA binding sites were downloaded from the Targetscan 6.2 database (21). To identify WES variants that were overlapping with these microRNA sites we used the BEDTools intersectBED function (44).

### Cloning of constructs for functional assays

To generate the microRNA expression constructs for miR-215, -342 and -346, regions encoding the primary transcripts were PCR amplified using the primers listed in Table S3 ('miRNAs'). The PCR products were then cloned using AgeI and EcoRI restriction sites in the pLKO.1 expression vector (Invitrogen) and the sequences were confirmed by Sanger sequencing.

miR-433 was initially cloned in the same way, but this construct did not express functional miR-433 when tested in cells (data not shown). As such an alternative approach was undertaken. MicroRNA primary transcripts have been successfully used to drive expression of synthetic short hairpin RNA (shRNA) to facilitate gene knock down (45). In such hybrid constructs, the sequence of the shRNA replaces the sequence that creates the stem-loop of the endogenous miR. Because we knew that miR-342 was successfully expressed (Figure S2), we inserted the stem-loop of miR-433 (which is responsible for generating the mature miR-433 sequence) into a plasmid already containing an expression cassette for miR-342. To facilitate removal of the miR-342 stem-loop from the miR-342 expression vector unique XbaI and SalI restriction sites were engineered at the 5' and 3' (respectively) of the miR-342 stem-loop. The miR-433 stem loop was amplified using primers containing XbaI and SalI restriction sites (detailed in Table S3) and the PCR product was cloned in place of the miR-342 stem-loop using XbaI and SalI. The sequence was confirmed via Sanger sequencing. Expression and functionality of this mir-433 construct was confirmed in a 'positive control' reporter assay (see below; Figure S2-A).

'Positive control' reporter constructs were used to confirm that all cloned microRNAs were expressed and able to regulate gene expression by interacting with ideal target sites in a reporter assay (Figure S2).Positive control reporters were generated as described previously (46, 47). Briefly, oligonucleotides containing 2 high-sensitivity binding sites for the cognate miRNAs were designed. To clone the reporter cassettes, sense and antisense oligonucleotides were generated and annealed to each other to form overhangs compatible with KflI restriction sites - allowing directional cloning into compatible sites. Oligonucleotides used for the generation of the reporters are listed in Table S3 ('miRNA positive control reporters').

The 3'UTR regions for each candidate gene spanning the patient identified variants (approximately 300-400 bp; see Figure 3A) were cloned into the pGL4.24 luciferase expression vector (Promega) as reporter constructs. Control human genomic DNA (Novagen human gDNA Cat. #69237) was used to PCR amplify the regions of interest (using the primers in Table S1; '3'UTR'). The sequences were confirmed via Sanger sequencing and shown to contain the control/reference allele. PCR products were cloned downstream of the luciferase reporter gene using XbaI and FseI restriction sites. Vectors carrying the alternative allele or with a deletion of the microRNA binding site were generated using the QuickChange Site-Directed Mutagenesis kit (Stratagene) following the manufacturer's instructions and using the primers in Table S3 ('3'UTR SDM'). The presence of the desired changes was confirmed via Sanger sequencing.

### Cell culture and transfection

We performed the reporter assay in HEK293 cells. HEK293 cells are a suitable cellular model as they are easy to culture and reach very high transfection efficiency. The protein machinery that microRNAs use to affect gene expression is ubiquitous and we have previously shown that this machinery is functional in HEK293 cells (46). All the experiments were carried out using HEK293 cells grown in DMEM (Invitrogen) media supplemented with 10% Fetal Calf Serum (Sigma) and 2mM Penicillin/Streptomycin. Cells were kept for the entire length of the experiments at 37 °C in presence of 5% CO_2_. Transfections were performed using GeneJuice (Novagen) following the manufacturer's instructions.

### Luciferase assay

3.0 × 10^4^ HEK293 cells were seeded in each well of a 24 well plate (60-70% confluence) 24 hours before transfection. Reporter constructs were co-transfected into cells alongside the microRNA expression vector and a Renilla reporter (pRL-TK) for internal normalization. 48 hours post-transfection, firefly luciferase and Renilla luciferase activities were measured as per manufacturer's instructions (Dual Luciferase reporter assay system, Promega).

### Websites

Exome Aggregation Consortium (ExAC), Cambridge, MA (URL: http://exac.broadinstitute.org) [last accessed January 2016]

The Genotype-Tissue Expression project portal (GTEx) (URL: http://www.gtexportal.org) [last accessed March 2016]

NCBI dbSNP Build 146 (URL: http://www.ncbi.nlm.nih.gov/SNP/) [last accessed January 2016] Targetscan Human (Prediction of microRNA targets) Release 6.2 (URL: http://www.targetscan.org/vert_61/) [last accessed January 2015]

miRBase, database of published miRNAs sequences and annotation (URL: http://www.mirbase.org/) [last accessed January 2015]

## ACKNOWLEDGMENTS

This work was funded by a Marie Curie Career Integration Grant and by a Max Planck Research Group Grant both awarded to S.C.V. The work of the Newbury lab is funded by the Medical Research Council [G1000569/1 and MR/J003719/1]. The UK Medical Research Council and the Wellcome Trust (Grant ref: 102215/2/13/2) and the University of Bristol provide core support for ALSPAC. We are extremely grateful to all the families who took part in the ALSPAC study, the midwives for their help in recruiting them, and the whole ALSPAC team, which includes interviewers, computer and laboratory technicians, clerical workers, research scientists, volunteers, managers, receptionists and nurses. This publication is the work of the authors and they will serve as guarantors for the contents of this paper. The work of the Wellcome Trust Centre in Oxford is supported by the Wellcome Trust [090532/Z/09/Z]. JH was supported by a scholarship from the Agency for Science, Technology, and Research, Singapore. We wish to thank Erik G. Willcutt, Richard K. Olson, John C. DeFries, Bruce F. Pennington and Shelley D. Smith for making available the CLDRC dataset.

## ABBREVIATIONS

NGS: Next generation sequencing
WES: Whole exome sequencing
WGS: Whole genome sequencing
SNV: Single nucleotide variant
UTR: Untranslated region
SLI: Specific language impairment
SNP: Single nucleotide polymorphism
miR: MicroRNA gene
kb: kilo bases
bp: base pairs
SLIC: Specific language impairment consortium
ELS: Expressive language score
RLS: Receptive language score
NWR: Non-word repetition
VIQ: Verbal intelligence quotient
PIQ: Performance intelligence quotient
CELF-R: Clinical Evaluations of Language Fundamentals
WORD: Wechsler Objective Reading Dimensions
MAF: Minor allele frequency
DEL: Deletion
eQTL: Expression quantitative trait loci
ALSPAC: Avon Longitudinal Study of Parents and Children
CLDRC: Colorado Learning Disabilities Research Centre
GWAS: Genome wide association study
shRNA: Short hairpin RNA
SDM: Site directed mutagenesis

